# Precocious reproduction in the ctenophore *Mnemiopsis leidyi* is modulated by conspecific density and food availability

**DOI:** 10.64898/2026.10.09.757825

**Authors:** Joan J. Soto-Angel, Elizaveta Ershova-Menze, Luis Martell, Pawel Burkhardt

## Abstract

Precocious reproduction, prior to the acquisition of the final adult morphology, occurs in a variety of animals and represents a distinct life-history strategy with important consequences for growth and survival. Larval reproduction is far rarer and usually restricted to groups that go through metamorphosis. In ctenophores, Carl Chun described “dissogeny” more than 130 years ago, a distinctive life-history pattern involving larval and adult reproductive periods separated by gonadal regression. Yet, the potential plasticity of larval reproduction in ctenophores remains unresolved. Here, we developed a method to follow and quantify reproductive output in *Mnemiopsis leidyi* larvae and tested environmental factors that may regulate precocious reproduction. We found that conspecific density strongly influences its onset: low-density, well-fed animals can bypass reproduction at the cydippid stage and develop rapidly into the lobed form, whereas higher density promotes precocious reproduction. Food availability and associated growth rate also contribute but cannot alone explain the observed differences. Once reproduction is established, however, high conspecific density suppresses reproductive output, an effect reproduced by exposure to seawater incubated with high *M. leidyi* densities, consistent with water-borne chemical communication. Salinity and light conditions further modulate spawning. Together, our results reveal substantial environmental plasticity in precocious reproduction in *M. leidyi*, reconcile long-standing contradictory observations of ctenophore development, and highlight how environmental cues can reshape reproductive life-history transitions.

## Introduction

Ctenophores, commonly known as comb jellies, are increasingly used to investigate fundamental questions in development, neurobiology, evolution, ecology and biogeography. Their phylogenetic position near the base of the animal tree makes them particularly informative for studying the origin and diversification of animal traits. Among them, *Mnemiopsis leidyi* has become one of the best-studied species, supported by genomic and single-cell resources (Ryan et al. 2013; Kim et al. 2025; Koutsouveli et al. 2025; Sebé-Pedrós et al. 2018) and by considerable interest in its ecological impact as a highly invasive species (Shiganova and Bulgakova 2000; Kideys 2002; Oguz et al. 2008; Piccardi et al. 2025). Its invasive success has been associated with high predatory efficiency, reproductive capacity, tolerance to starvation, regenerative ability and pronounced developmental plasticity (Colin et al. 2014; Jaspers et al. 2011; Granhag and Hosia 2015; Jaspers et al. 2015; Ramon-Mateu et al. 2019; Soto-Angel and Burkhardt, 2024). Yet several fundamental aspects of its reproductive biology remain poorly understood. This knowledge gap partly reflects broader challenges of ctenophore research: ctenophores are exceptionally fragile and difficult to maintain under laboratory conditions, and studies of their reproductive biology have therefore been limited and often rely on wild-caught adults (Courtney et al. 2020; Patry et al. 2020; Presnell et al. 2022; Soto-Angel et al. 2024; Sasson and Ryan 2016; Sasson et al. 2018).

One particularly enigmatic feature is precocious, or larval, reproduction. This phenomenon was first documented in ctenophores by Carl Chun in 1892. Chun studied the lobate species *Bolinopsis vitrea* and *Leucothea multicornis* and introduced the concept of dissogeny as an entirely new mode of reproduction in animals: two periods of reproduction (larval and adult) separated by a period of gonadal regression. Similar reproductive strategies were later reported in several ctenophore lineages, including *Pleurobrachia bachei* (Garbe 1901; Totton 1954; Hirota 1972), *Pleurobrachia globosa* (Wang et al. 2020), *Mertensia ovum* (Jaspers et al. 2012), *M. leidyi* (Martindale 1987; Edgar et al. 2022) and *Beroe* sp. (Edgar et al. 2022). However, other studies did not detect precocious reproduction in ctenophores (Baker and Reeve 1974; Kremer 1976), leading some authors to question whether dissogeny exist at all (cf. Reeve and Walter 1978). More recently, Edgar et al (2022) reported continuous reproduction beginning shortly after hatching and proposed that precocious reproduction is an intrinsic feature of ctenophore development. Under this interpretation, ctenophores would be direct developers and concepts such as larva and dissogeny would require reconsideration.

These apparently contradictory observations raise a fundamental question: is larval reproduction expressed by default, or is it a plastic life-history trait that can be environmentally regulated? Chun himself suggested that reproductive larvae might originate from similarly precociously reproductive individuals, hinting at flexibility in the life cycle. Yet the environmental factors governing the onset and maintenance of precocious reproduction have not been systematically tested. This question is particularly relevant for *M. leidyi*, a simultaneous hermaphrodite with two conspicuous life-history phases: a tentaculate cydippid stage, also known as larva, and a lobed stage, referred to as adult (Soto-Angel et al. 2024). Here, we developed a reproducible culture system and a quantitative assay to follow reproductive output in *M. leidyi* during the cydippid larval stage. We tested whether precocious reproduction can be promoted or suppressed by ecologically relevant factors, focusing particularly on conspecific density and food availability, and examined how physicochemical and biological conditions modulate reproductive output once reproduction is established. This framework allowed us to distinguish between the capacity for precocious reproduction and the environmental regulation of its expression.

## Results

### Establishment and validation of experimental design

#### A stable culture based on larval reproduction

Consistent with previous observations (Edgar et al. 2022), *M. leidyi* can begin reproducing at the cydippid stage within days of hatching. Under our culture conditions, reproduction was observed as early as six days after hatching and typically began after 8-10 days. Depending on the culture conditions, cydippids could continue growing while reproducing through and beyond lobe development (Fig. 1). Alternatively, cydippids could be maintained at a body length of 3-4 mm while reproducing continuously for at least 2.5 years under appropriate feeding, water-exchange and densities conditions. One main culture unit (MCU) containing 8-10 larvae, 3-4 mm oral-aboral (OA) length in 200 mL of seawater and fed daily with *Brachionus* produced several dozen offspring per day. The system could be scaled to produce several thousands of larvae per week with relatively few resources. For example, 20 beakers yielded more than 4000 larvae per week, while requiring less than 25 L of seawater, including the rotifer culture. By contrast, low densities (ca. 1-4 animals per MCU) combined with high feeding levels promoted rapid development, with animals developing lobes within approximately 10-14 days after hatching and, in some cases bypassing reproduction at cydippid stage. At this stage (ca. 1 cm), these animals could then be transferred to larger containers and grown to 8-10 cm.

**Figure 1.**
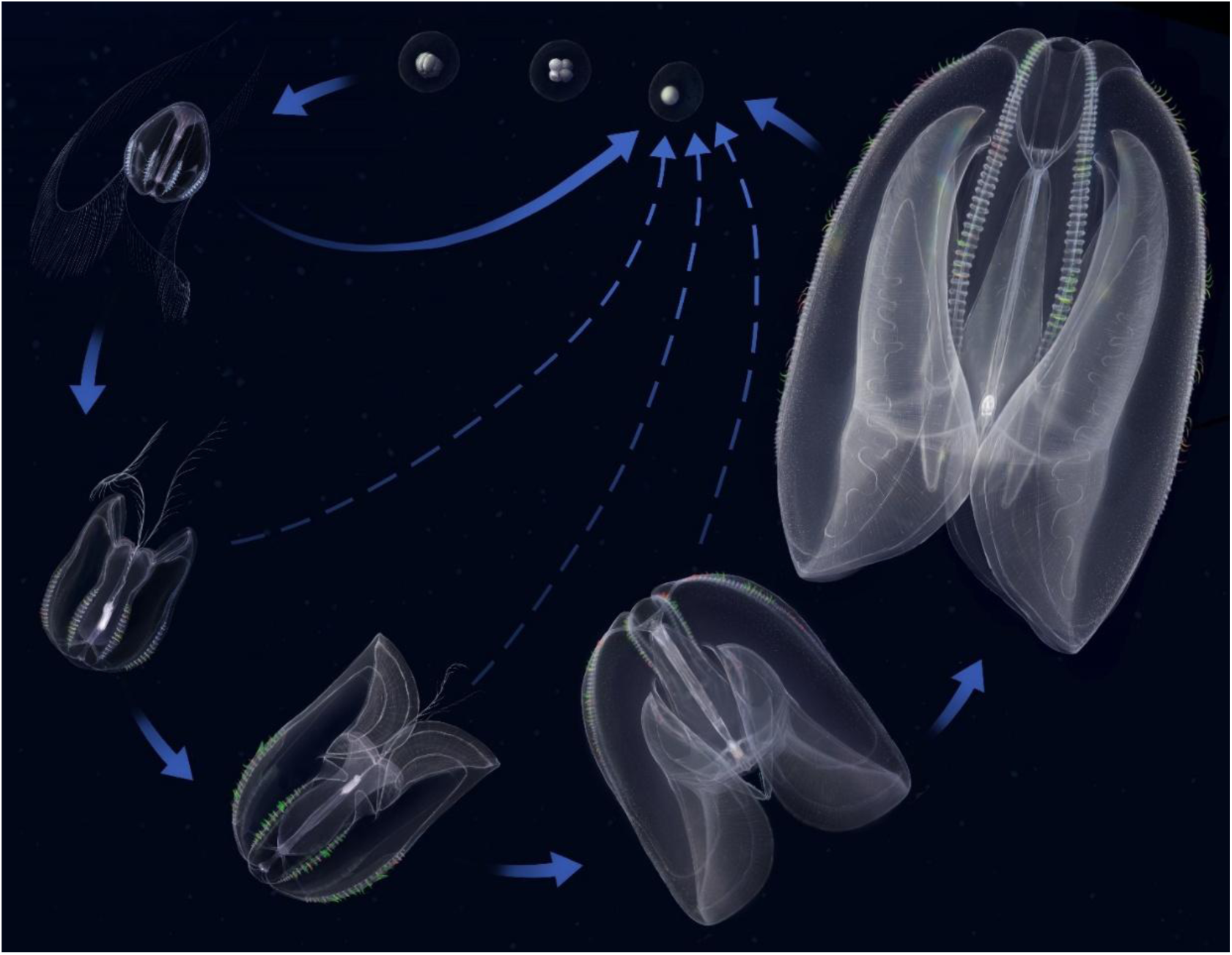
Life cycle of *Mnemiopsis leidyi*. The lobate ctenophore *Mnemiopsis leidyi* is a simultaneous hermaphrodite with external fertilization and two main life-history phases: the cydippid and the lobed stages. Eggs are fertilized shortly after spawning and larvae hatches approximately 24 hours after fertilization. The cydippid larva is characterized by the presence of two long and retractable tentacles used to capture prey. Within a couple of weeks after hatching, animals enter a transition phase and begin developing new feeding structures, first the lobes and subsequently the auricles. Concurrently, the tentacles are reabsorbed while the lobes and auricles continue to grow. Large lobed adults can reach approximately 10 cm in total length. Strikingly, *M. leidyi* can start reproducing as early as six days after hatching and continue reproducing during subsequent development. Illustration by Nick Bezio.

#### Clearance rate defines feeding requirements for experimental conditions

The rotifer *Brachionus* provides a suitable prey source for *M. leidyi* larvae. Its size (approximately 90-150 µm) is suitable for recently hatched cydippid larvae, and it can be cultured and used as the sole food source to maintain healthy, continuously reproducing larvae. Rotifers have previously been used to successfully raise *M. leidyi* larvae (Presnell et al. 2022) and support precocious reproduction (Edgar et al. 2022). Defining an appropriate feeding level was therefore important both for maintaining larvae at a stable size and for establishing a quantitative baseline for experiments in which food availability was manipulated.

Gut content increased rapidly during the first 20 minutes after feeding and subsequently stabilized at approximately 17 ± 5 prey items per gut (Fig. 2A). On the contrary, digestion time slowly increased over the same period, from approximately 30 ± 4 minutes during the first 10 min to 55 ± 18 minutes after 60 min (Fig. 2B), when the guts had the highest amount of prey items. Clearance rate peaked 20 min after feeding and decreased thereafter (Fig. 2C). The overall mean clearance rate was 2.85 ± 1.0 ml per individual and hour (n=42), with a median of 2.80 ml ind^-1^ h^-1^ (n=42) closely matching the value after 60 min (2.87 ± 0.7 ml ind^-1^ h^-1^; n=12). Combining gut content, digestion time and prey concentration allowed us to estimate clearance rates under our culture conditions, with values consistent with previous findings using single larvae (Finenko et al. 2008; Sullivan 2010). Based on these rates, 10 larvae in 200 ml of seawater with a prey density of 8.5 prey*/*ml would be expected to deplete the available prey in approximately 7 hours: In practice, larvae 3-4 mm in OA length remained at a stable size over time when fed with 180-200 *Brachionus* per individual per day, corresponding to 9-10 prey/ml in the MCU and approximately one third of their estimated clearance capacity. Lower feeding levels generally resulted in shrinkage, whereas feeding levels approaching the estimated clearance capacity promoted growth.

**Figure 2.**
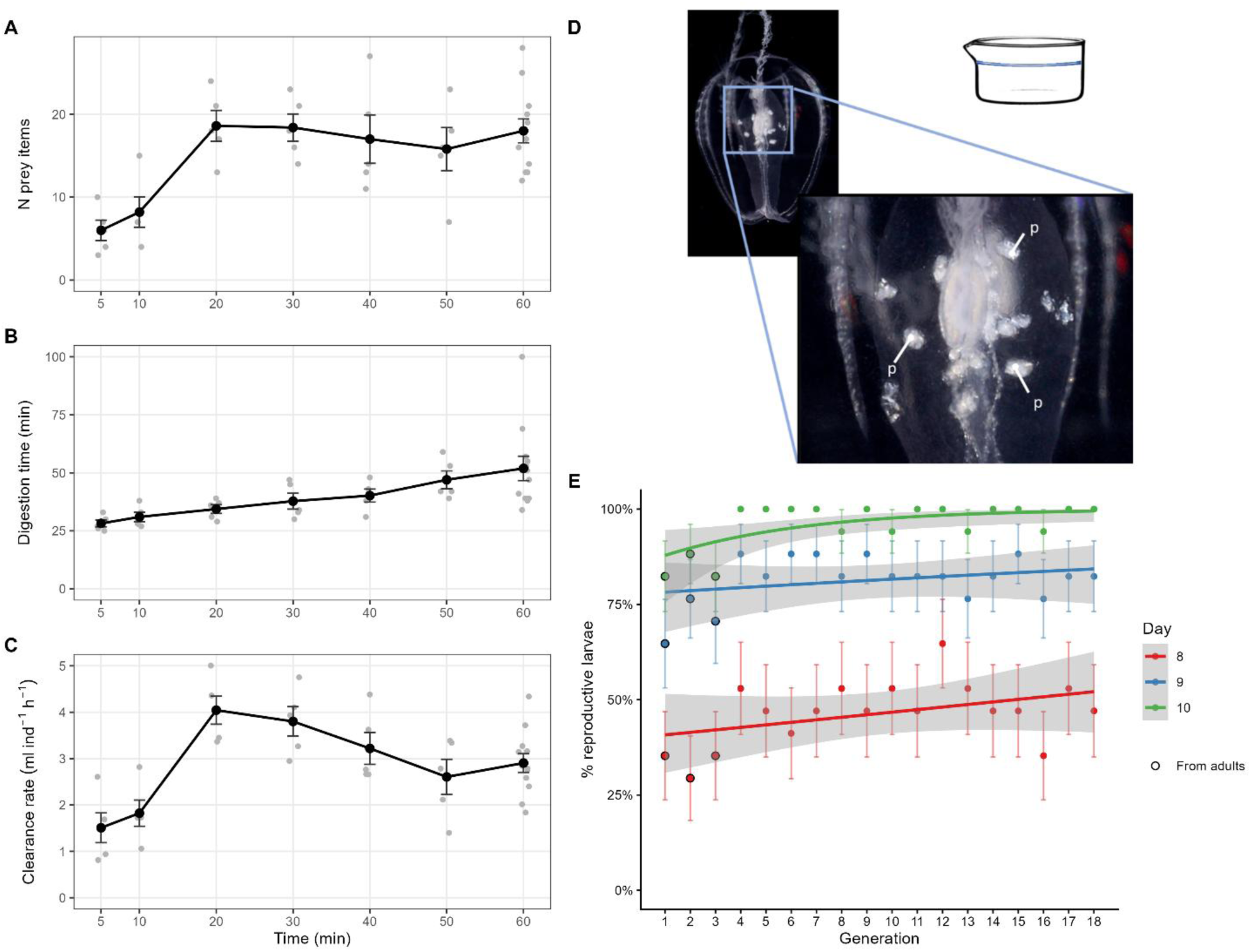
Experimental validation of feeding conditions and reproductive stability. A-C. Feeding dynamics of *Mnemiopsis leidyi* larvae over 60 minutes: (A) Mean number of prey items per larvae over time; (B) Digestion time over the same time. (C) Clearance rate, expressed as ml per individual and hour. Black points show means ± SE and grey points individual larvae. (D) *M. leidyi* larva containing several rotifers in the gut (p). The transparency of *M. leidyi* allows prey items in the gut to be quantified, enabling real-time monitoring of prey capture and digestion. (E) Proportion of larvae reproductive by days 8, 9 and 10 across 18 consecutive generations. Points show proportions ± SE; lines show binomial GLM fits with 95% confidence bands. By day 10, 96% of all larvae were reproductive, and more than 80% in every generation.

#### Other potential experimental limitations: cannibalism and inbreeding

We found no direct or indirect evidence of cannibalism during the cydippid stage. Despite the high density of potential prey, none of the 3-5 mm larvae assessed contained eggs or recently hatched larvae in the gut in the first experiment, nor remnants of *Brachionus* previously ingested by the recently hatched larvae in the second experiment. In addition, direct observations showed that recently hatched larvae were not entrapped by the tentacles or tentilla of larger animals, but rather passed over them. By contrast, we occasionally observed eggs and/or larvae in the guts of small lobed *M. leidyi* of approximately 15-30 mm in OA length, consistent with previous reports of cannibalism in larger animals (Javidpour et al. 2020). Together, these observations suggest that cannibalism is absent or negligible during the cydippid stage. We therefore did not apply correction factors or additional measures to account for larval predation in experiments using with 3- 5 mm larvae.

Across 18 consecutive generations, the proportion of reproductive larvae did not change significantly with generation on days 8 or 9, but increased slightly by day 10 (Binomial GLM; odds ratio per generation 1.21, p = 0.008). However, this effect disappeared when only larvae obtained from other larvae (generations 4-18) were included in the model. By day 10 after hatching, 96% of all larvae were reproductive, prior to lobe development (Fig. 2E). All larvae reproduced at least once during the observation period. Similarly, 65% of the larvae obtained from large wild adults spawned by day 10, with all of them spawning by day 15 (n=119). Interestingly, wild-caught 2–4 mm cydippids also retained the capacity to reproduce, although reproduction was delayed in many individuals after collection. Specifically, 3 out 11 larvae spawned on day 5 after collection, 2 between day 6-10, and the remaining 6 only after at least 12 days. These results indicate that all *M. leidyi* larvae have the potential –physiological machinery– to reproduce during the cydippid stage. Hence, we assume no inbreeding effect in this regard for our culture (i.e. the Baltic population used for single-cell sequencing, Sebé-Pedros et al. 2018). Nevertheless, all experiments were performed using animals from the same generation to minimize potential generational effects and variability.

### Factors triggering larval reproduction in recently hatched larvae

Recently hatched larvae raised at different densities substantially diverged in size measured on day 10. Mean body size decreased with increasing density, revealing an inverse relationship between density and growth (Fig. 3A) corresponding to a decrease of 0.77 mm per doubling of density (Linear mixed model; 95% CI 0.60- 0.94 mm, p < 0.001). No spawning was observed in any MCU containing 2 larvae per 200 ml, and 4 of 5 MCUs containing 4 larvae per 200 ml also showed no spawning during the 14-day experiment. By contrast, spawning was observed from day 9 onward in the highest-density MCUs (8 and 16 larvae per 200 ml), and by day 13 all MCUs at these densities contained reproductive larvae. Firth logistic regression predicted an odds ratio of 25 per doubling of density (95% CI 1.0–618, p = 0.048) (Figure 5). These differences were unlikely to result from competition for food because prey abundance was adjusted proportionally to larval density, keeping food availability per larva constant. Furthermore, food was supplied at levels sufficient to maintain prey availability throughout the experiment. Likewise, monitored water-quality parameters showed no deterioration in the high-density treatments that could explain the observed differences (see methods section). Thus, increasing density was associated with an increased probability of precocious reproduction but reduced somatic growth. Because animals were not followed individually, we cannot determine whether density affected growth and reproduction independently, or whether density triggered reproduction, which in turn reduced growth rate.

**Figure 3.**
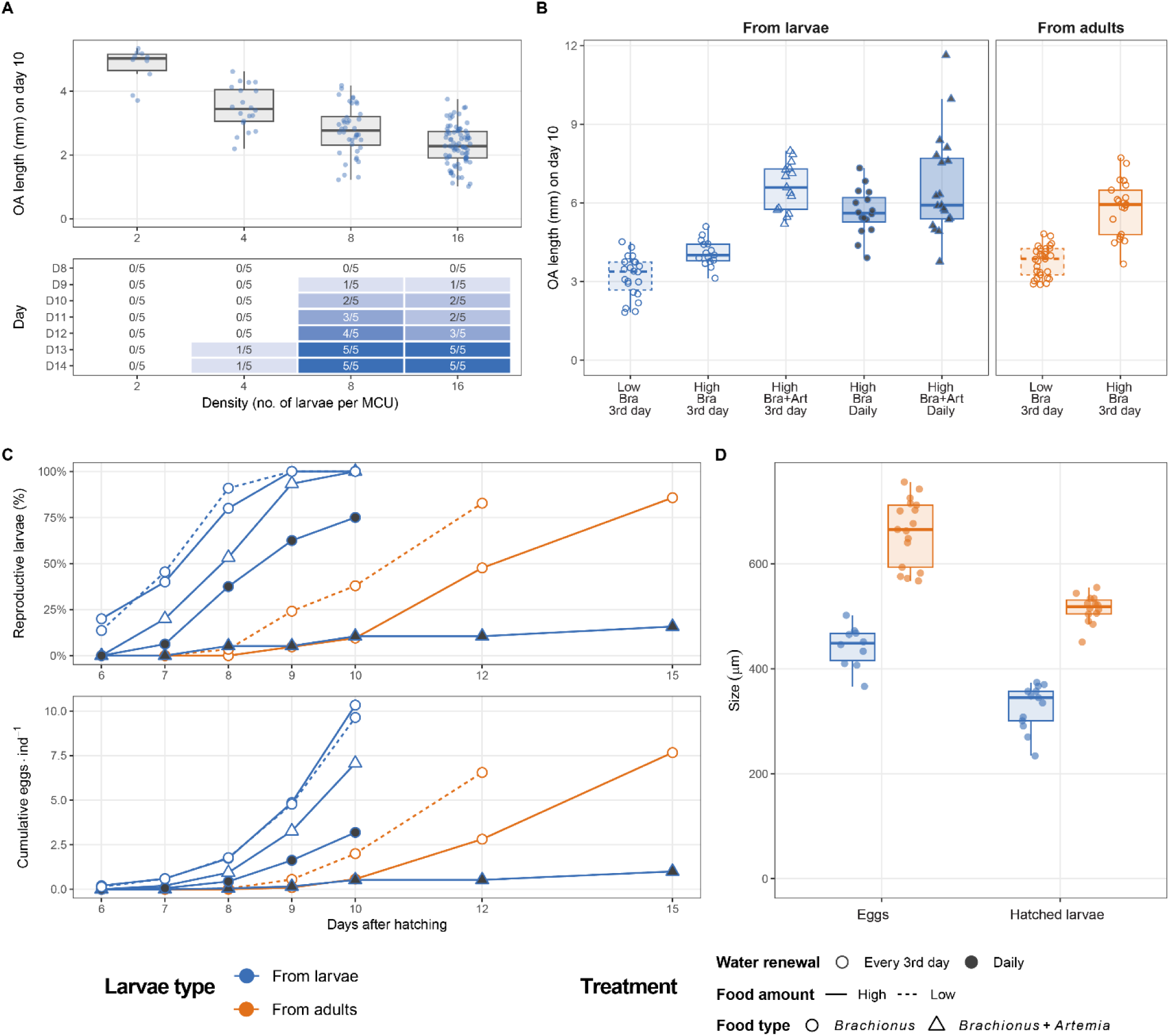
Factors triggering larval reproduction in recently hatched larvae. (A) Effects of density on body growth and on the onset of precocious reproduction. Top: oral-aboral (OA) length on day 10 of larvae reared at densities of 2, 4, 8 or 16 larvae per 200 ml. Box plots show the median and interquartile range, and points represent individual larvae. Bottom: number of MCUs in which spawning was observed from days 8-14; shading indicates the proportion of MCUs showing spawning at each density and time point. (B) OA length on day 10 of the individually reared offspring produced by reproductive larvae (blue) or wild adults (orange) under different combinations of food amount (high, low), food type (Bra = Brachionus, Bra+Art = Brachionus + Artemia) and frequency of water renewal (daily or every third day). Box plots show the median and interquartile range. (C) Cumulative proportion of reproductive larvae over time (top) and mean egg production per individual for each treatment (bottom). D. Comparison of egg and hatchling size between offspring produced by reproductive larvae and adults. Box plots show the median and interquartile range, and points represents individual eggs or larvae.

We next tested additional factors using individually monitored larvae, allowing reproductive onset, body size and reproductive output to be followed for each animal. The rate of water exchange and food type availability were also identified as significant triggers in larval reproduction, timing of maturation and reproductive output. When water was exchanged every third day, food amount significantly affected body size on day 10, with high food producing 31% larger larvae than low food (GLS; ratio of geometric means = 1.31, CI 1.15-1.48, p < 0.001) (Fig. 3B), whereas the timing of first reproduction and overall reproductive output were nearly identical between feeding levels (p = 0.79 and 0.46, respectively) (Fig. 3D). Among offspring of reproductive larvae fed exclusively with *Brachionus* and water exchanged every third day, all animals had reproduced by day 9. By contrast, daily water renewal with the same feeding conditions produced 62% of reproductive larvae on day 9 and 75% by day 10 (Fig. 3C). Adding *Artemia* to the diet delayed the onset of reproduction by approximately one day relative to larvae fed exclusively with *Brachionus*, although this difference was not statistically significant (Cox PH test; HR=0.54, 0.26-1.1, p = 0.18). Daily water renewal delayed reproduction both in larvae fed only *Brachionus* (Cox PH test; HR 0.21, 0.10-0.48) and those with a diet combining *Brachionus* and *Artemia* (HR=0.005, 0.001-0.02; both p < 0.001), and reduced egg production by 69% and 93%, respectively (RR=0.31, 0.17-0.56, and 0.07, 0.03-0.17; both p < 0.001). Noteworthy, the strongest effect was observed under high feeding with both *Artemia* and *Brachionus* combined with daily water exchange, where only 2 of 19 (11%) animals were reproductive by day 10. Daily water renewal increased the growth in larvae fed only *Brachionus* (GLS; ratio of geometric means = 1.38, 1.24-1.52, p < 0.001), but not in larvae with the Artemia-enriched diet (0.98, 0.28-1.13, p = 0.79).

Overall, the largest larvae occurred in treatments combining high food availability with frequent water exchange. However, their mean size did not differ significantly from larvae fed with *Brachinous* and exposed to daily water exchange or from larvae fed both prey types in water exchanged every three days (Fig. 3B). The effect of daily water renewal on maturation remained after accounting for larval length (Cox PH test; *Brachionus*: HR 0.40, 0.16-0.95, p = 0.039; *Brachionus* + *Artemia*: HR 0.018, 0.003-0.09, p < 0.001). Within treatments, larger larvae matured later (Cox PH; HR 0.70 per mm, 0.52-0.95, p = 0.023), although, as length was measured after the onset of reproduction, the direction of this relationship cannot be resolved. Most larvae receiving both prey type had well-developed lobes by day 10, but only those exposed to daily water renewal commonly bypassed reproduction at the cydippid stage. By day 15, all 19 animals in this treatment had developed into small lobed ctenophores of approximately 1 cm OA length, yet only 3 were reproductive. Mean egg production per individual, including animals that did not spawn, followed the same overall pattern: daily water exchange was associated with low or absent egg production and greater somatic growth (Fig. 3B, 3C).

Larvae obtained from large wild animals reached a larger size by day 10 than those produced by larvae under the same feeding and water renewal regimes, both at high (GLS; ratio of geometric means 1.41, 1.27-1.56, p < 0.001) and low (1.2, 1.06-1.36, p = 0.011) food amounts. (Fig. 3B). This can be explained by the larger size of the eggs and larvae obtained by large adults versus the ones produced by 3-4 mm larvae, which were 0.22 and 0.26 mm larger, respectively (Welch t-test; p < 0.001) (Fig. 3D). Despite their larger size, most larvae from large wild adults reproduced later than those from larvae under the same food and water-renewal conditions (Cox PH; low food: HR 0.05, 0.02-0.11; high food: 0.03, 0.01-0.07, p < 0.001) and produced fewer eggs by day 10 (RR 0.21, 0.13-0.35; 0.06, 0.03-0.12, respectively, both p < 0.001). Most offspring of large wild adults under low-food treatment did not reproduce until day 12, whereas under high-food treatment reproduction was delayed until approximately day 15. Under equivalent feeding and water-exchange regimes, first reproduction therefore occurred approximately 3–5 days later than in offspring of reproductive cydippids (Fig. 3C). High food amount further delayed reproduction (Cox PH; HR 0.40, 0.20-0.82, p = 0.035) and reduced egg production (NB GLM; RR 0.29, 0.14-0.60, p = 0.003) compared with low food. Some offspring of adults exposed to high food-amount, had already developed lobes before reproducing for the first time. Among offspring of reproductive cydippids, this rapid growth was observed only under daily water exchange (Fig. 3C).

Together, these experiments show that conspecific density and water-exchange frequency strongly influence the onset of precocious reproduction. Our findings confirm a clear link between precocious reproduction and growth rate, as previously documented by Martindale (1987). In particular, animals subjected to frequent water renewal generally grew faster and reproduced later or bypassed larval reproduction, whereas animals under less frequent water exchange remained smaller and reproduced earlier. However, because some rapidly growing individuals nevertheless reproduced by day 10, growth rate alone cannot explain precocious reproduction occurs (Fig. 5). Offspring of large adults also tended to reproduce later than offspring of reproductive cydippids, indicating that developmental starting conditions may additionally influence reproductive timing.

### Factors modulating reproductive output in established reproductive larvae

For larvae maintained for several months under stable conditions and reproducing continuously, changes in salinity had pronounced effects on the reproductive output. Reducing salinity from 34 to 28 ppt increased reproductive output to an average of 2.4 fold above the control (RR 2.36, 95% CI 1.38–4.03, p < 0.001) (Fig. 4A). Conversely, increasing salinity from 34 to 37 ppt strongly inhibited reproduction, with approximately fourfold fewer offspring than in controls maintained at constant salinity (RR 0.29, 0.16–0.54, p < 0.001). A smaller decrease of 2 ppt produced no significant change in reproductive output (RR 1.39, 0.81–2.40, p = 0.34). Interestingly, following a decrease in salinity, reproductive output approached control levels (RR 1.36, 0.79–2.34, p = 0.41). By contrast, inhibition persisted during the second spawning event following an increase in salinity (RR 0.24, 0.13–0.45, p < 0.001). The stimulatory effect of reduced salinity was reproduced across a second salinity range (Fig. 4B). At 26 ppt, output doubled relative to both the initial value (RR 2.15, 95% CI 1.59–2.92) and the control (RR 2.13, 1.50–3.04; both p < 0.001). After the return to 30 ppt, it fell to less than half of the initial value (RR 0.45, 0.32–0.63) and of the control (RR 0.47, 0.32–0.68; both p < 0.001). Output in the control did not change over time (p ≥ 0.23) (Fig. 4B).

**Figure 4.**
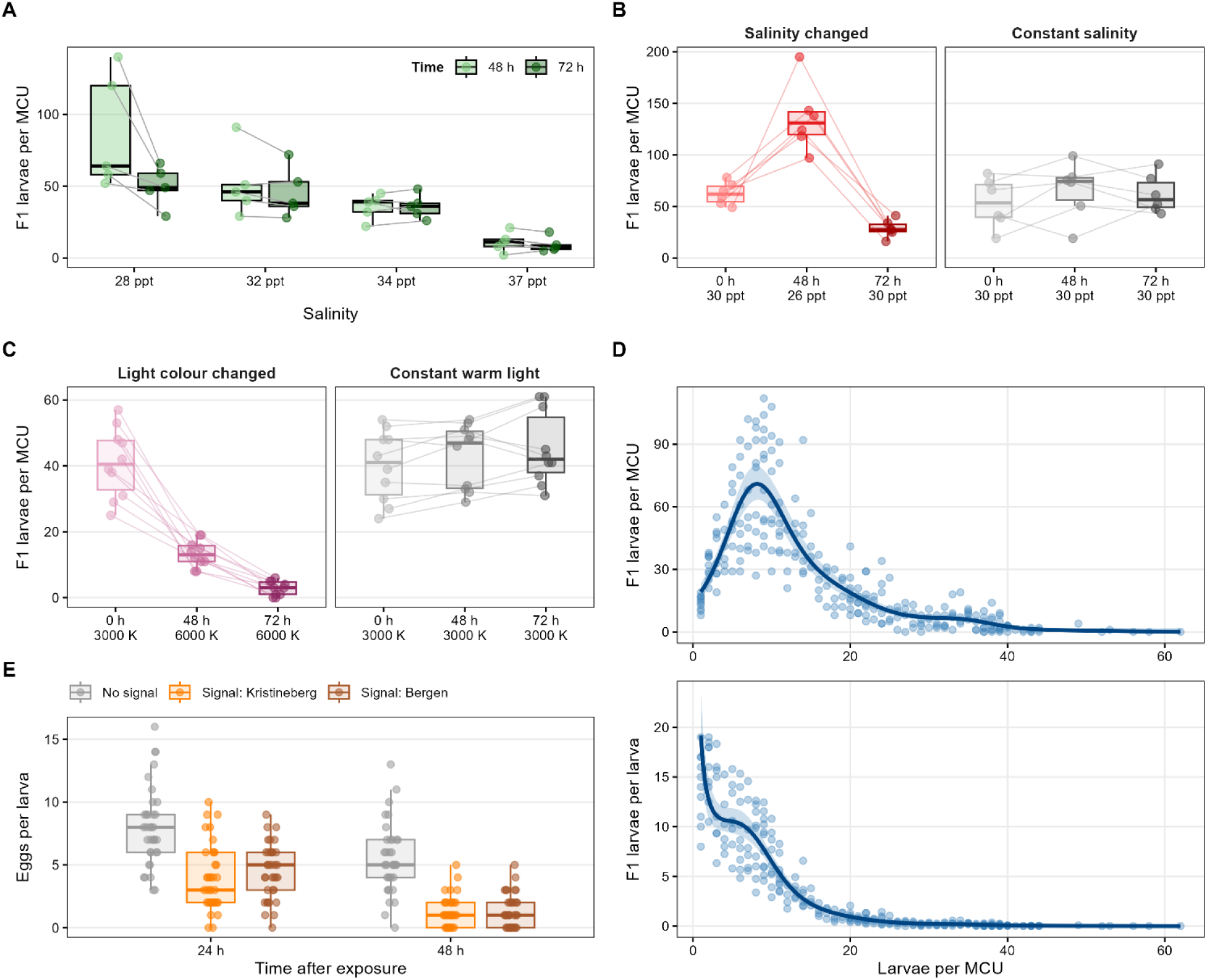
Environmental modulation of reproductive output in established reproductive larvae. (A-C) Effect of salinity (A-B) and light color temperature (C) on reproductive output across successive spawning events. Box plots show the median and interquartile range; points represent individual MCUs, and thin lines connect the same MCU across spawning events (D) Effect of conspecific density on reproductive output per MCU (above) and per individual (below). Solid lines and shaded bands show the fit of a negative binomial generalized additive model with 95% confidence intervals. (E) Spawning inhibition through exposure to high densities in successive spawning events. Individual larvae were maintained in control seawater (“No signal”) or in seawater conditioned for 24 h by high densities of conspecific larvae originated from Kristineberg (Skagerrak) or Bergen (North Sea). Box plots show the median and interquartile range; points represent individual larvae. Grey indicates controls, and lighter to darker shades of each colour indicate successive spawning events.

Changing light color temperature also strongly affected reproductive output. Switching from warm (3000 K) to cold (6000 K) illumination reduced the number of eggs produced by reproductive larvae to 31% in the first spawning event (RR 0.31, p < 0.001) and to 6% in the second spawning event (RR 0.06, p < 0.001) (Fig. 4C). Larvae maintained continuously under cold illumination likewise showed low reproductive output, comparable to that observed during the second spawning event.

Reproductive output also varied non-linearly with conspecific density (nb GAM, 80% deviance explained, p < 0.001). MCUs containing 8 individuals produced the highest total number of offspring per culture (Fig. 4D). By contrast, MCUs containing more than 30 animals rarely produced more 15 offspring per day. When reproductive output was expressed per individual rather than per MCU, increasing density was associated with a pronounced decline in offspring production per animal. Thus, offspring production per larva decreased rapidly as the number of reproductive animals per MCU increased (Fig. 4D). To test whether this effect could be mediated by a water-borne conspecific factor rather than by direct crowding or associated changes in feeding conditions, we exposed individual larvae to seawater previously conditioned by high densities of conspecifics (see methods section). During the first spawning event, larvae exposed to conditioned seawater spawned roughly 50% less eggs than controls (Kristineberg: RR 0.49; Bergen: RR 0.57, both p < 0.001) (Fig. 4E). The inhibitory effect was even stronger during the second spawning event (Kristineberg: RR 0.24; Bergen: RR 0.25, both p < 0.001). Conditioned seawater produced by animals from Kristineberg Marine Station (Skagerrak) and Bergen (North Sea) had comparable effects (p = 0.79).

## Discussion

### Is larval reproduction an intrinsic feature of the ctenophore life cycle?

Precocious reproduction in ctenophores has been debated since its original description over 130 years ago (Chun 1892). The discrepancies among previous studies have led to several, sometimes mutually exclusive interpretations. Dissogeny –two periods of reproduction separated by a period of gonad regression– was originally described by Chun for two lobate species (*Bolinopsis vitrea* and *Leucothea multicornis*) and later supported by Garbe (1901), Totton (1954) and Wang et al. (2020) for two cydippids (*Pleurobrachia bachei* and *Pleurobrachia globosa*), by Martindale (1987) for *M. leidyi*, and by Jaspers et al. (2012) for *Mertensia ovum*. Conversely, other authors did not find any evidence of precocious reproduction in *M. leidyi* (Baker and Reeve 1974; Kremer 1976) or even suggested that dissogeny did not occur in ctenophores (Reeve and Walter 1978). More recently, Edgar et al. (2022) reported continuous reproduction in *M. leidyi* beginning early after hatching, as well as early reproduction in *Beroe* sp. The authors concluded that precocious reproduction is an intrinsic part of the ctenophore life cycle based on its occurrence in distantly related species across the ctenophore phylogeny.

Here we confirm that *M. leidyi* can reproduce continuously from as early as 6 days after hatching and beyond the proposed pause in reproduction (Martindale 1987), in agreement with Edgar et al. (2022). Importantly, however, our experiments show that expression of this capacity is not obligatory: under some environmental conditions, *M. leidyi* bypasses reproduction at the cydippid stage altogether. Precocious reproduction therefore appears to represent an intrinsic capacity of *M. leidyi* that can be modulated, rather than a fixed developmental programme that is invariably expressed (Fig. 5). By contrast, several attempts to induce larval reproduction in both cultured and wild-caught *Bolinopsis infundibulum* under the same density and feeding conditions used for *M. leidyi*, were unsuccessful, suggesting that this species either lacks the capacity for larval reproduction or requires different environmental triggers. Whether larval reproductive competence is broadly shared across ctenophores therefore remains unresolved. For most ctenophore species, early life- cycle stages and larval reproductive competence remain fundamentally unexplored, underscoring how incomplete our understanding of ctenophore life cycles remains (Soto-Angel et al. 2023).

**Figure 5.**
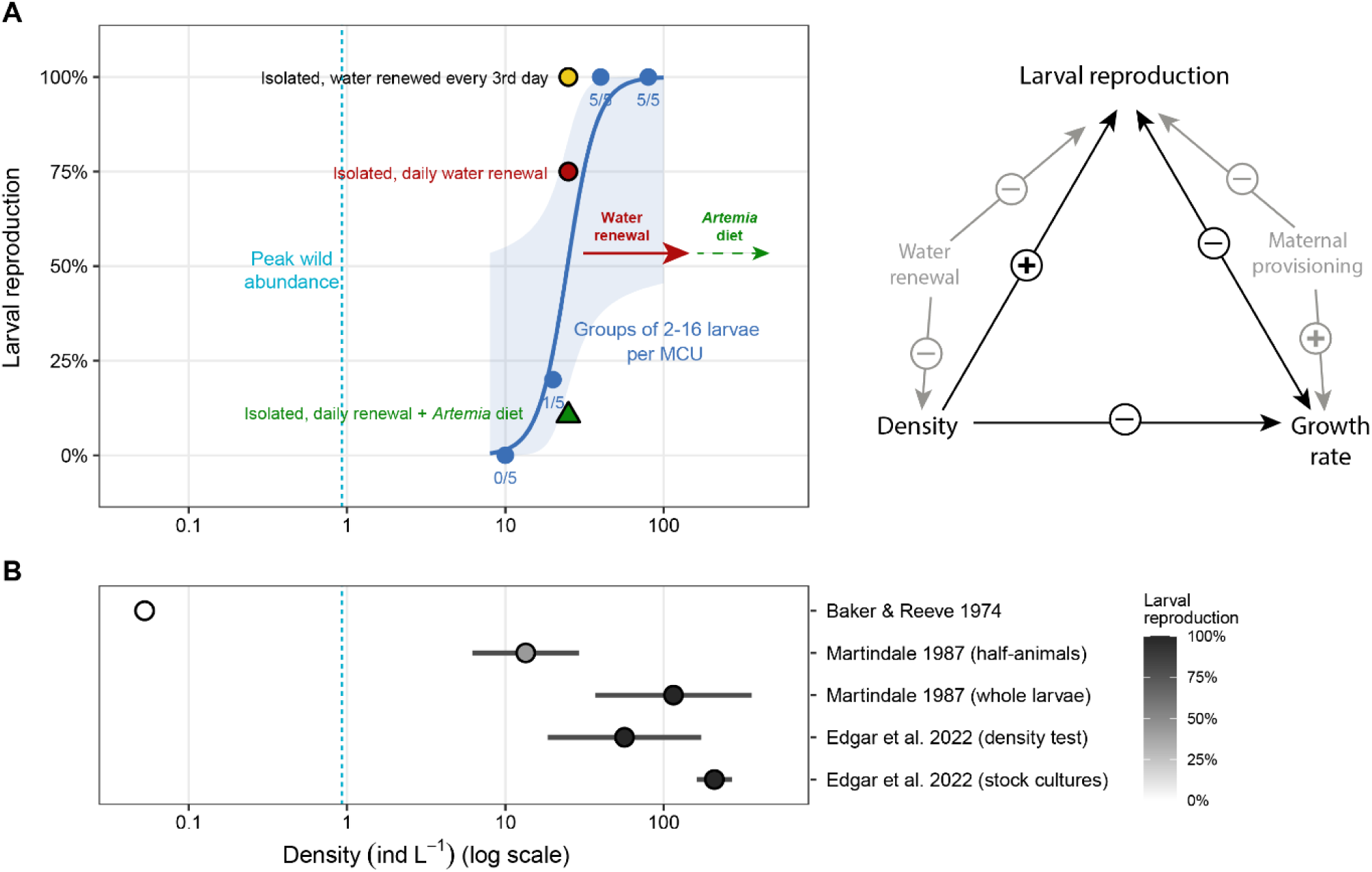
Density dependence of precocious reproduction in *Mnemiopsis leidyi*. (A) Relationship between culture density and precocious reproduction in the present study. Blue symbols represent groups containing 2-16 larvae per MCU and indicate the proportion of MCUs in which spawning was observed by day 14. The curve and shaded band show a Firth-penalized logistic regression on log₁₀-transformed density and its 95% confidence interval. Additional symbols show the effects of other experimental treatments, and arrows show their approximate direction relative to the density-response relationship. (B) Comparison with previous studies of precocious reproduction in *M. leidyi*. Horizontal ranges indicate the approximate culture densities used in each study, with shading representing the reported occurrence of precocious reproduction. The dashed line indicates the approximate maximum abundance reported in the field for small *M. leidyi* (Riisgård et al. 2007).

The present study documents larval reproduction in populations of *M. leidyi* outside their native range, consistent with observations from Mediterranean specimens (Macarena Marambio pers. comm). Earlier maturation of lobed *M. leidyi* in invasive compared to native populations has been attributed to variability in reproductive strategies, on which selection can act during the invasion process (Jaspers et al., 2017). We observed that already the first generation derived from wild-caught animals reproduced within a few days after hatching, with no consistent change in the proportion of reproductive larvae across 18 laboratory generations after day 10. This indicates that larval reproductive competence is maintained over long-term culture rather than representing a transient response to recent collection from the wild.

Among the environmental variables tested, conspecific density emerged as a major determinant of whether this reproductive capacity was expressed. Specifically, at low densities, *M. leidyi* skips reproduction at the cydippid stage even when food is abundant. This density dependence provides a potential explanation for the differences observed by previous authors. Baker and Reeve (1974) raised *M. leidyi* individually in 19 L tanks and observed reproduction only when individuals reached ca. 3 cm. In contrast, Martindale (1987) concentrated the larvae in 75 to 300 ml beakers and described a dissogenic reproductive pattern, with first reproduction at the cydippid stage at 1.8-2.8 mm, and the second when the animals were 35 mm in size. Both authors used natural zooplankton as a diet, but the animals raised by Baker and Reeve (1974) at lower densities had a substantially higher growth rate (approximately 10 times higher than in Martindale (1987) and ca. 4 times higher than the values reported here). More recently, Edgar et al. (2022) reported continuous reproduction along cydippid, transitional and lobed stages. Thus, studies of the same species under significantly different culture conditions produced markedly different results.

The divergent results reported to date, and the interpretations drawn from them, are likely influenced by experimental design, which may not reflect conditions in the wild. Ctenophore fecundity, for instance, is known to depend strongly on experimental conditions such as container volume (Purcell et al. 2001). Here, even the lowest density values used are above the highest abundances of small *M. leidyi* reported in the wild: 867 ± 121 individuals m^-^³ (Riisgård et al. 2007). For larger adults, abundances of 0.2 to 0.4 individuals m^-3^ have been frequently recorded under bloom conditions, with local values reaching 2 animals m^-3^ (Hosia and Falkenhaug 2015). Edgar et al. (2022) performed their density experiments in small 50 ml containers with 1- 8 larvae each and enriched rotifers as a diet. Our results show that even a single animal in 50 ml can reproduce precociously under these conditions. Only when seawater is renewed daily or animals are maintained at densities below 20 larvae per litre, is the proportion of animals displaying larval reproduction significantly reduced. As we also used enriched *Brachionus* fed with the same commercial brand of microalgae (RGcomplete), the experimental designs differed primarily in the densities tested, which may contribute to contrasting conclusions regarding the effect of density on cydippid reproduction. Altogether, these comparisons suggest that responses observed under high laboratory densities may not fully reflect conditions experienced in the wild, except perhaps during dense bloom events (Fig. 5).

### Density and growth as cues for precocious reproduction

Density-dependent modulation of reproduction is known across several animal groups. In rotifers such as *Brachionus*, high density triggers a switch from asexual to sexual reproduction via a self-released chemical cue (Stelzer and Snell, 2003). The nematode *Caenorhabditis elegans* integrates population-density cues to adjust reproductive timing and duration (Wong et al., 2020). Among vertebrates, some salamander species use larval density as a cue to switch between early paedomorphic reproduction and reproduction delayed until after metamorphosis (Harris 1987; Doyle and Whiteman, 2008). Across these systems, population density can provide information about local environmental conditions, including habitat stability, resource availability and competition (Cohen, 1966; Pianka, 1970). High density may therefore act as a composite cue rather than reflecting a single environmental variable (Wilbur and Collins, 1973). In *Amblystoma* species, unstable environments such as short-lived ponds favour relatively early metamorphosis, whereas permanent ponds permit long larval periods and the appearance of sexually mature paedomorphs (Harris 1987; Doyle and Whiteman, 2008).

Our results show that *M. leidyi* skips larval reproduction in conditions of low density combined with high food availability (Fig. 5), conditions consistent with a stable, resource-rich environment with limited intraspecific competition. Under these conditions, *M. leidyi* larvae may experience less pressure for precocious reproduction, with investment instead shifting toward somatic growth and, consequently, greater reproductive output at larger body size (Baker and Reeve, 1974; Sasson and Ryan, 2016). Conversely, high conspecific density may favour precocious reproduction in environments with high competition or predation pressure. In ctenophores, an extreme case is known for *Mertensia ovum* in the central Baltic Sea, where individuals reproduce exclusively through larval reproduction, and do not reach the final adult morphology; a strategy interpreted as favouring precocious reproduction under high predation risk (Jaspers et al. 2012). *Mnemiopsis leidyi* is a bloom-forming species that has recently been reported to cannibalize larvae (Javidpour et al. 2020), which may further contribute to the risks associated with high-density *M. leidyi* blooms beyond competition for resources. Reproductive timing in *M. leidyi* larvae appears highly plastic rather than fixed: early reproduction is favoured under high-density conditions and suppressed when resources are abundant and conspecific density is low.

Chun (1892) already noted that eggs and larvae produced by other larvae were comparatively smaller than those produced by large lobed animals and hypothesized that precociously reproductive individuals originated from other reproductive larvae. One possible explanation is differential maternal provisioning: larger, structurally mature adults may produce larger eggs than those from precociously reproducing larvae, resulting in offspring that begin development at a larger size (Marshall and Keough 2008). For *M. leidyi*, a larger starting size coupled with adequate food availability may promote faster growth and delayed reproduction, thereby favouring the bypass of precocious reproduction (Fig. 5). The offspring produced by larvae remain comparatively smaller by the time they reach reproductive competence. The delayed reproduction observed in the offspring of large adults may therefore be a developmental consequence of this size advantage rather than an independently transmitted trait, consistent with reports of parental investment shaping offspring life-history trajectories in other taxa (Reichert et al. 2019) and providing a plausible proximate mechanism for Chun’s original observations.

Hirota (1972) and Martindale (1987) noted that precociously reproductive larvae grow more slowly than non-reproductive ones but could not determine the direction of the cause–consequence relationship. Overall, our results are consistent with this pattern, with larger animals preferentially skipping reproduction at the larval stage. However, we show that growth rate alone cannot account for the occurrence of larval reproduction: manipulating feeding regimes had little effect on reproduction rates when the density was kept unchanged, despite considerable differences in body size on day 10 (Fig. 3). This argues against reduced growth being sufficient to induce precocious reproduction, although it does not resolve whether slower growth is partly a cause or a consequence of reproduction. Smaller individuals preferentially allocate resources to growth over gamete production (Sasson and Ryan 2016), which could contribute to the association between body size, growth and reproductive state. Comparative data between offspring of adults and larvae under low water renewal conditions point in the opposite direction: low water renewal (and thus higher effective density) was not linked with precocious reproduction, leaving open the possibility that growth or starting body size also modulates reproductive timing (Fig. 5). Similar challenges in disentangling the effect of density and growth rate are known in amphibians (Harris 1987), with at least one example supporting a density effect on precocious reproduction independent of growth rate (Richter et al. 2009).

### Factors modulating reproductive output in established reproductive larvae

Reproductive output in adult *M. leidyi* appears to be shaped by multiple, interacting factors rather than a single driver. Temperature sets a lower threshold below which reproduction does not occur, but once this threshold is exceeded, output becomes highly variable and is not consistently explained by temperature alone (Sobrero et al. 2026). This variability has been attributed to additional factors, including body size, food availability, and salinity (Baker and Reeve 1974; Finenko et al. 2006; Jaspers et al. 2011; 2015; Shiganova 2020; Rečnik et al. 2024). Among these, salinity has received particular attention, though its effect appears most pronounced toward the lower end of the species’ tolerance range. In the Baltic Sea, where salinity is markedly reduced relative to the species’ native range, experimental work has shown that reduced salinity directly limits reproduction and constrains population expansion (Jaspers et al. 2011), consistent with reports that egg production and larval development cease below a salinity of ∼6–6.5 ppt in the Caspian and Azov Seas (Shiganova 2020). In the northern Adriatic, blooming populations have been recorded across a broad salinity range (11–38 ppt) with high measured fecundity under high-salinity conditions (Malej et al. 2017). Rather than exposing animals to constant salinity treatments across a gradient, here we assessed the response to a salinity shift. Our results show that *M. leidyi* larvae modulate their reproduction in response to a change in salinity, with the direction of that change determining whether reproductive output is stimulated or inhibited. A shift to higher salinity can physiologically stress the animals, sometimes causing damage or even disintegration (Yazdani Foshtomi et al. 2007; Soto-Angel et al. 2024), which may account for the inhibitory effect observed here following an increase in salinity. In contrast, a decrease in salinity may indicate exposure to more estuarine conditions, where shallow, productive areas provide particularly favourable conditions for *M. leidyi* (Siapatis et al. 2008), consistent with its native distribution in estuaries along the Atlantic coast (Purcell et al. 2001). The stimulatory effect of reduced salinity observed here may therefore reflect sensitivity to environmental changes associated with favourable estuarine conditions. Whether this sensitivity to a shift in salinity is specific to larval reproduction or also applies to the adult stages remains to be tested.

*Mnemiopsis leidyi* possesses opsin genes capable of detecting light, with at least one opsin’s absorption spectrum overlapping its own bioluminescent emission (Schnitzler et al. 2012), providing a plausible basis for light sensitivity. Behavioral evidence supports this capacity: adults discriminate between wavelengths, reacting rapidly to blue light but little to red (Silakov et al. 2023). Reproductive output itself has been considered largely independent of light: spawning is synchronized with the light–dark cycle but its timing is governed primarily by an endogenous circadian rhythm rather than by an immediate response to light or darkness (Sasson and Ryan 2016). Likewise, experimental UV exposure affected growth and survival but not egg production (Rathjen et al. 2012). Here, we show that a shift from warm to cold light colour considerably reduces reproductive output. Whether this response reflects an ecologically relevant spectral signal remains to be tested.

Here, we demonstrate that conspecific density has stage-dependent effects on larval reproduction: high densities promote precocious reproduction in recently hatched, pre-competent larvae once a critical threshold is exceeded, but suppress reproductive output in several-month-old, post-competent larvae with established reproductive activity. Population density has been shown to impose differential stage-dependent effects on both reproductive timing and output in other taxa (Mugabo et al. 2013; Müller et al. 2016). In *M. leidyi*, precocious reproduction may reflect elevated risk during blooms, when prey depletion can drive adult conspecifics to cannibalize larvae (Javidpour et al. 2020), consistent with elevated mortality risk promoting earlier maturation and reproductive effort (Reznick et al. 1990). By contrast, competent larvae could benefit from delaying spawning until density declines and competition eases. These results suggest that conspecific density acts as a stage-dependent environmental cue, promoting the onset of precocious reproduction in recently hatched animals while suppressing reproductive output once a stable reproductive state has been reached. Conspecific communication is known to regulate spawning in adult *M. leidyi* (Sasson et al. 2018), and to delay maturation and reduce offspring number in other planktonic invertebrates (Carvajal-Salamanca et al. 2008). Our findings provide evidence that spawning in *M. leidyi* larvae can be regulated by conspecific cues, opening the way to investigate the identity and mechanism of the underlying signal or signals.

Taken together, our results help reconcile more than a century of conflicting observations on precocious reproduction in ctenophores. In *M. leidyi*, precocious reproduction appears to represent an intrinsic but highly plastic reproductive capacity, whose expression is strongly influenced by conspecific density and other environmental conditions. This plasticity highlights the remarkable flexibility of the *M. leidyi* life cycle and provides a framework for understanding how reproductive timing may be adjusted to changing ecological conditions.

## Methods

### Preconditions for experimental design and methodology validation

#### 1. Establishing a culture based on larval reproduction

The first generation of reproductive *Mnemiopsis leidyi* cydippids was obtained from adults collected at Kristineberg Marine Research Station (Gothenburg, Sweden) in Skagerrak. Well-fed adults were allowed to spawn in 2 L beakers kept in complete darkness overnight. The following day, the eggs were pipetted individually and transferred to 300 mL beakers (ca. 300 eggs/beaker) previously incubated in seawater for a minimum of 24 h to prevent egg adhesion to the bottom. Salinity and temperature were kept constant during all transfers. More details on the spawning procedure for adults can be found in Soto-Angel et al. (2024). Once the eggs hatched (ca. 22-26 h after spawning), the cydippid larvae were pipetted and transferred to the main culture unit (MCU). The MCU consists of a 300 mL crystallizing dish with a spout, 95 mm wide and 55 mm high, covered with the base of a tissue culture dish with approximately the same outer diameter as the inner diameter of the crystallizing dish. Each MCU was filled with 200 mL of seawater and populated with 50- 100 recently hatched larvae, which were fed daily with ca. 4000-5000 *Brachionus* (rotifers) per MCU during the first 5 days. On day 5, the culture was split, reducing the density to 15-20 animals/MCU. At this stage, each MCU is fed with 2500-3000 *Brachionus* per day. On day 8, the culture was split again and the density reduced to 8-10 animals/MCU. Each MCU was then fed with ca. 2000 *Brachionus* per day (5 times a week). The seawater was filtered through 10, 5 and 1 µm meshes, irradiated with UV, purified with activated charcoal, and diluted with reverse-osmosis water to a final salinity of 27 ± 0.5 ppt and pH 7.8-8.1 (normal values unless otherwise specified). Animals were kept at constant room temperature (21 ± 1 °C under a constant photoperiod (9 h light, 15 h dark). MCUs were placed on a transparent surface with fluorescent lamps underneath (colour temperature 3000 K). For culture performance, see the Results section.

#### 2. Feeding standardization

To standardize the feeding regime and ensure its reproducibility, we estimated the potential food intake of larvae 3-4 mm in oral-aboral (OA) length fed with the rotifer *Brachionus*. Clearance rate was calculated as the volume cleared per animal and unit time, based on prey concentration, gut content, and digestion time (Granhag et al. 2011):

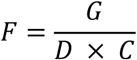

where F: Clearance rates (L ind^-1^ h^-1^), G: Gut content (prey ind^-1^), D: Digestion time (h), C: Prey concentration (prey L^-1^).

Because single time-point measurements may not capture temporal variation in feeding and digestions, gut content and digestion time were assessed at 5, 10, 20, 30, 40, 50, and 60 min after feeding, while individual prey items were still identifiable in the gut. Eight MCUs (MCU1, MCU2…), each with 10 larvae, 3-4 mm in OA length were used, matching the standard larvae density and size in our culture conditions. Each MCU was fed with ca. 1700±110 *Brachionus*, resulting in an initial concentration of 8.5 prey/mL. Five minutes after feeding, 5 larvae were randomly collected from MCU1 and the number of prey items in their guts (i.e. digestion rate) was counted under a stereomicroscope. The same procedure was followed 10 minutes after feeding for MCU2, 20 minutes after feeding for MCU3, and at subsequent time points up to 60 mins. Digestion time, defined as the time until gut contents were completely digested, was recorded for all larvae surveyed. An additional set of larvae from MCU8 was examined for gut content and digestion after time 60 min to further capture potential interindividual variability. Prey concentration was corrected at each time point based on the mean number of prey items removed and total number of larvae per MCU.

The mean clearance rate was used to estimate the maximum number of rotifers provided in subsequent experiments. Unless otherwise specified, MCUs were fed with ca. 2000 *Brachionus* per day (5 days per week). The amount of *Brachionus* was estimated using a cell counter prior to the experiments and adjusted to the required value if necessary. More details on larval feeding and maintenance of the rotifer culture can be found in Soto-Angel et al. (2024).

#### 3. Constant water quality throughout the experiment

In some cases, the experimental design included treatments with high feeding levels (i.e., values above the clearance rate) or high densities of animals in relatively small-volume containers. Prior to these experiments, we determined the maximum values of food concentration and larval density that could be maintained in an MCU without compromising the water quality. In addition, water quality was monitored before, during and after the different experiments. If significant variations occurred, the experiment was terminated and the results were disregarded. Salinity, pH, temperature, dissolved oxygen, and nitrogen compounds (ammonium, nitrites, and nitrates) were used as indicators of water quality. Salinity and temperature variations within 5% of the starting value were considered acceptable. For pH, variations within 2% were considered satisfactory. Any detectable concentration above 0 was used as the cutoff for nitrogen compounds. For dissolved oxygen, saturation above 85% and concentrations above 8 mg/L, together with no significant differences between treatments or experiments were considered adequate.

#### 4. Quantification of reproductive output in *M. leidyi* larvae

Different methods for quantifying reproductive output in adult *Mnemiopsis leidyi* include reverse filtration using a slow flow through a fine mesh (cf. Jaspers et al. 2014) or estimating the total number of eggs by sampling an aliquot of the egg-containing volume and extrapolating to the total volume (Sasson and Ryan 2016). These methods allow rapid quantification of large numbers of eggs. However, unlike large adult *M. leidyi*, which can produce thousands of eggs per day (Baker and Reeve, 1974), cydippid larvae rarely produce more than 20 eggs per individual per day (Martindale 1987). Therefore, to increase accuracy and reduce uncertainty, eggs were counted manually after being pipetted from the MCU.

Reproductive output in larvae was quantified by transferring the animals to new MCUs containing clean seawater and allowing them spawn overnight. *M. leidyi* larvae spawn daily; the eggs are fertilized shortly after being shed and they hatch approximately 24 h after fertilization at 20 °C. Hence, two spawning events can be monitored simultaneously: 48 h after the reproductive animals were transferred, the MCU contained the eggs produced during the preceding 24 h, and the larvae that had hatched from eggs produced 24-48 h earlier. The eggs and recently hatched larvae were readily identified using a transmitted light source.

#### 5. Occurrence of larval reproduction in wild animals and through different generations

To assess potential effects of inbreeding in the *M. leidyi* culture, a multigenerational experiment was performed. Eighteen MCUs were populated with 100 recently hatched larvae each. Each MCU contained a different generation (F1-F18), with the first three generations (F1, F2 and F3) produced by adults ca. 5 cm in OA length. All MCUs were raised in the same conditions and simultaneously (i.e. standard water parameters and feeding regime as described above). On day 5, a subset of 17 randomly selected larvae was transferred individually to 50 ml crystallizing dishes and fed with 400-500 *Brachionus* per day per beaker. The dishes were carefully monitored for signs of spawning from days 7-14 after hatching, when larvae can already display larval reproduction (cf. Martindale 1987; Edgar et al. 2022).

To evaluate potential differences in the proportion of precocious larvae between laboratory-raised and wild-caught animals, as well as between closely related lobate species *Bolinopsis infundibulum*, the same procedure was applied for larvae obtained from seven large (ca. 10 cm) *M. leidyi* adults and four large *B. infundibulum* (ca. 13 cm) collected from several localities around Bergen. The larvae obtained from these animals were raised under the same conditions described above, isolated in 50 ml crystallizing dishes on day 5, and checked for signs of spawning from days 8-14 after hatching. In addition, several wild *M. leidyi* and *B. infundibulum* larvae, 2-4 mm in OA length, were sampled and checked for signs of spawning immediately after collection and during the following two weeks. For *Bolinopsis infundibulum* the latter experiment was repeated again on board RV Heincke during HE605 Arctic expedition RISING in Svalbard. Animals (n=33) were kept at ca. 2 °C, fed with *Brachionus* and followed for 10 days for signs of spawning. Only healthy and intact larvae were included in the experiment.

#### 6. Cannibalism in larval *M. leidyi*

*Mnemiopsis leidyi* adults have been reported to prey on *M. leidyi* cydippid larvae (Javidpour et al. 2020), which could potentially bias estimates of reproductive output if the density of the parental generation was high enough to substantially reduce the number of eggs and/or larvae present in the container. To test whether cannibalism also occurs at cydippid stage, ten larvae 3-5 mm OA length, starved for 24 h before the experiment, were transferred to a single MCU with over 1500 eggs and recently hatched larvae. The gut contents of the 3-5 mm larvae were examined under a stereomicroscope after 20 min, 1 h and 3 h, with the animals returned to the MCU between observations. To evaluate whether the ingested larvae were rapidly digested, thereby preventing their identification in the gut, a second experiment was performed. In this experiment, the recently hatched larvae were previously fed with *Brachionus*, so that rotifer remains could serve as an indicator of larval ingestion. In addition to the time periods mentioned, 15 min direct observations were conducted every hour.

### Factors inducing or inhibiting gonad development in recently hatched larvae

#### 1. Effect of density on growth rate and larval reproduction

Recently hatched larvae were placed in MCUs at different densities, with 2, 4, 8 and 16 larvae per MCU and five replicates per density. MCUs were fed proportionally to the number of larvae per unit, providing 300 prey items per animal per day for the first 5 days, and 600 rotifers per animal per day thereafter. Animals were transferred to new MCUs containing renewed seawater every 3 days to maximize water quality. Room temperature and standard salinity and pH values were used (see culture section). The MCUs were checked daily for signs of spawning, and the larval OA length was measured on day 10. Measurements were taken using Adobe Photoshop measurement tool on calibrated photographs acquired with a Canon 5D Mark IV and Canon MP-E 65 mm macro lens. The experiment was stopped on day 14.

#### 2. Effect of food regime and density in the timing and intensity of larval reproduction

Based on the results, a second experiment was designed to test additional variables and quantify the timing and reproductive output at the individual level. Recently hatched larvae were placed individually in 50 ml crystallizing dishes and fed progressively over the course of the experiment. Three factors were assessed: food amount, food type and water renewal. Two levels were tested for food amount. Under the low-food regime, animals were fed progressively with 2-8 *Brachionus*/ml (i.e. 80-320 *Brachionus* per larva). Under the high-food regime, 10-15 *Brachionus*/ml (i.e. 400-600 *Brachionus* per larva) were used. Food type was tested only under the high-food regime. Two food types were considered: *Brachionus* alone and *Brachionus* + *Artemia*. *Artemia* was added progressively after day 5, with 0.5-2 nauplii/ml (i.e. 20 to 80 nauplii per larva). Given that *Artemia* nauplii are too large to be efficiently captured and ingested by *M. leidyi* larva only a few days old (which is nearly the same size as the recently hatched *Artemia*), no food regime was based exclusively on *Artemia*. Water renewal was also tested only under the high-food regime, with two levels: daily and every three days. In the latter case, larvae were pipetted in the same manner but returned to the same dish, thereby controlling for the effects of handling and transfer. The frequency of water renewal was used to manipulate the accumulation of potential water-borne cues, allowing the potential effects of conspecific accumulation on the timing and intensity of larval reproduction to be tested at the individual level. Larvae were kept in the containers for 12-15 days, and total length (including lobes when present) were measured on day 10 following the procedure described above. The day of first reproduction was recorded, and the number of eggs produced per animal was counted daily.

#### 3. Differences between offspring of adults and reproductive larvae

To compare potential differences between offspring produced by larvae and by adults at the individual level, offspring obtained from three large adults were raised as described above under the low- and high-food regimes, with *Brachionus* as the only food type. The larvae were kept in the containers for 15 days and were equally measured for total length (including lobes) on day 10 using the same procedure described above. The day of first reproduction was recorded, and the number of eggs produced per animal was counted daily. In addition, a selection of eggs and recently hatched larvae produced by large adults and by reproductive larvae were photographed and measured under the microscope using Leica software. Eggs were measured twice, with the measurement axis rotated by 90° for the second measurement, and the average of both measurements was calculated. For recently hatched larvae, OA length was measured. These eggs and early larvae were not used for further experimentation.

### Factors modulating reproductive output in already-reproducing larvae

In the following experiments, 2- to 3-month–old larvae, 3-4 mm in OA length, were exposed to changes in salinity, light colour, and density. All larvae used were reproductively active and spawning daily at the start of each experiment. The reproductive output was measured under control conditions (no changes) and following the respective treatment. Larval age and generation were kept constant within each experiment.

#### 1. Effect of salinity

A random selection of 200 well-fed larvae raised at 34 ppt and a density of 10 animals per MCU were transferred to new MCUs containing seawater at four different salinities: 28, 32, 34 and 37 ppt (10 larvae per MCU and five replicates per treatment). Two consecutive spawning events were monitored: 48 h after the animals were transferred, all hatched larvae present in the MCU (i.e. the first spawning event) were counted and removed; 24 h later (72 h after transfer), all the larvae present in the MCU were counted again (i.e., the second spawning event). To test whether the response was consistent across a different salinity range, a second experiment was conducted with larvae raised at 30 ppt instead. In this experiment, reproductive output was quantified both before and after transfer across two additional spawning events (six replicates for control and six replicates per treatment, 10 animals per MCU). The quantification of reproductive output was made as described for the first experiment.

#### 2. Effect of light colour temperature

Two hundred well-fed larvae raised at 34 ppt and a density of 10 animals per MCU were randomly selected and transferred to new MCUs containing seawater with the same water parameters. Ten MCUs with 10 animals each were maintained under standard culture conditions and exposed to a warm light source of 3000 K, simulating sunset/sunrise light conditions. The remaining 10 MCUs were simultaneously transferred to a different bench with the same light intensity but illuminated with a cool light of 6000 K, simulating daylight conditions. The same photoperiod (as described in the culture section) was used for both treatments. Two contiguous consecutive spawning events were monitored as described above.

#### 3. Effect of density

More then 4000 reproductive larvae were distributed in MCUs in different densities, ranging from 1 to 40 animals per MCU, with a minimum of four replicates per density level. The animals were maintained at these densities for 48 h and fed with *Brachionus* proportionally, at 300 *Brachionus* per larvae. After 48 h, the animals were carefully removed, and the seawater was filtered twice through 50 µm mesh to remove all the uneaten food and spawned offspring. The animals were moved back to their corresponding MCUs with the filtered seawater, and newly produced larvae were counted 48 h later (first spawning event).

Based on the results, a second experiment was designed to obtain individual-level data and exclude alternative explanations for the observed trends, such as metabolites or other by-products released by the prey. Twenty-four MCUs were filled with seawater, eight of them populated with 30 larvae each from our main culture (Kristineberg population), and another eight with 30 larvae each from the local Bergen population. All larvae were starved for 2 h before the experiment to allow gut evacuation and were not fed thereafter. The remaining eight MCUs were used to obtain control seawater and were treated identically except that no *M. leidyi* larvae were added. All MCUs were equally incubated at room temperature for 24 h, after which the seawater was filtered twice through a 50 µm mesh and divided into 50 ml crystallizing dishes. A different set of 107 well-fed larvae from our main culture (Kristineberg), previously maintained at a density of 10 animals per MCU, was transferred individually to 50 ml crystallizing dishes, with 37 replicates receiving seawater conditioned by Kristineberg larvae, 35 receiving seawater conditioned by Bergen larvae, and 35 receiving control seawater. After 24 h, all eggs present in the dishes were removed and counted (1^st^ spawning event). The procedure was repeated 24 h later (48 h after transfer; 2^nd^ spawning event).

### Statistical analysis

All analyses were performed in R 4.4.2 (R Core Team 2024). Binomial generalized linear models (GLMs) were used to calculate the probability of being reproductive on days 8, 9 and 10, with generation as a continuous predictor. The proportion of MCUs spawning by day 14 in the density experiment was modelled against log₁₀ density with Firth’s penalized logistic regression. Larval length on day 10 was analysed with a linear mixed model with log₂ (density) as fixed effect and MCU as a random factor using the package glmmTMB (Brooks et al. 2017). Whether spawning occurred in an MCU by day 14 was analysed with Firth’s penalized logistic regression on log₂(density) (Firth 1993), with a Cochran-Armitage test for trend. Length on day 10 of individually reared larvae was analysed with generalized least squares (GLS) on log-transformed length, with a separate variance for each treatment via the nlme package (Pinheiro et al 2024). Time to first reproduction was analysed via the package survival (Therneau 2024) using a Cox proportional-hazards model (Therneau and Grambsch 2000), with larvae right-censored on the last observation day of their treatment. Cumulative egg production up to day 10 was analysed with a negative binomial GLM (Venables and Ripley 2002). For length, maturation and egg production, eight planned contrasts (food amount, food type, water renewal and origin) were tested with Holm adjustment. The effect of water renewal on maturation independent of growth was tested with a Cox model including water renewal, food type, their interaction and length on day 10, fitted to high-food larvae from larvae. The association between length and maturation within treatments was tested with a Cox model stratified by treatment. Because length was measured after the onset of reproduction, length effects are reported as associations. Sizes of eggs and recently hatched larvae produced by adults and by larvae were compared with Welch’s t-tests. The number of F1 larvae per MCU was analysed with a negative binomial generalized linear mixed model (GLMM) with salinity, spawning event and their interaction as fixed effects and MCU as the random intercept. Each salinity was compared with the control (34 ppt) using Dunnett contrasts via the emmeans package (Lenth and Piaskowski 2026). The number of F1 larvae per MCU was analysed with negative binomial GLMMs with treatment, spawning event and their interaction as fixed effects and MCU as random intercept. The number of F1 larvae per MCU was modelled with a negative binomial generalized additive model (GAM) using the mgcv package (Wood 2011; 2017). Eggs per larva were analysed with a negative binomial GLMM with water treatment, spawning event and their interaction as fixed effects and larva as random intercept. Each conditioned-water treatment was compared with the control using Dunnett contrasts.

## Acknowledgments

The authors want to express their gratitude to Anne Aasjord, Eilen Myrvold, Alexandre Jan (Michael Sars Centre) and Kim Øvrebø (ILAB) for their support and advice on water quality assessment. We thank Aino Hosia (UiB), Sanna Majaneva (NTNU/Akvaplan-niva) and Cornelia Jaspers (DTU) for insightful discussions on life cycle plasticity and adaptations in ctenophores. Special thanks go to Nick Bezio for the detailed illustration of the *M. leidyi* life cycle. We would like to thank the crew of RV Heincke during the HE605 RISING expedition to Svalbard, with special acknowledgment to cruise leader Charlotte Havermans (AWI). This work was supported by the Michael Sars Centre core budget, through the Research Council of Norway (NFR project no. 234817) and the European Research Council Consolidator Grant (101044989, “ORIGINEURO”) awarded to P.B.

## References

Baker, L.D., Reeve, M.R. (1974) Laboratory culture of the lobate ctenophore *Mnemiopsis mccradyi* with notes on feeding and fecundity. Marine Biology 26(1):57–62. 10.1007/BF00389086

Brooks, M.E., Kristensen, K., van Benthem, K.J., Magnusson, A., Berg, C.W., Nielsen, A., Skaug, H.J., Maechler, M., Bolker, B.M. (2017) glmmTMB balances speed and flexibility among packages for zero-inflated generalized linear mixed modeling. The R Journal 9:378–400. 10.32614/RJ-2017-066

Carvajal-Salamanca, J.L., Aranguiz-Acuña, A., Ramos-Jiliberto, R., Zúñiga, L.R. (2008). Immediate and delayed life-history responses of Daphnia ambigua to conspecific cues. Journal of Plankton Research 30(10):1117–1122. 10.1093/plankt/fbn071

Chun, C. (1892) Die Dissogonie, eine neue Form der geschlechtlichen Zeugung. In: Festschrift zum siebzigsten Geburtstage Rudolf Leuckarts. Engelmann, Leipzig, pp 77–108

Cohen, D. (1966) Optimizing reproduction in a randomly varying environment. Journal of Theoretical Biology 12(1):119–129. 10.1016/0022-5193(66)90188-3

Colin, S.P., MacPherson, R., Gemmell, B., Costello, J.H., Sutherland, K., Jaspers, C. (2014) Elevating the predatory effect: sensory-scanning foraging strategy by the lobate ctenophore *Mnemiopsis leidyi*. Limnology and Oceanography 60(1):100–109. 10.1002/lno.10007

Courtney, A., Merces, G.O.T., Pickering, M. (2020) Characterising the behaviour of the ctenophore *Pleurobrachia pileus* in a laboratory aquaculture system. bioRxiv. 10.1101/2020.05.25.114744

Doyle, J.M., Whiteman, H.H. (2008) Paedomorphosis in *Ambystoma talpoideum*: effects of initial body size variation and density. Oecologia 156(1):87–94. 10.1007/s00442-008-0977-2

Edgar, A., Ponciano, J.M., Martindale, M.Q. (2022) Ctenophores are direct developers that reproduce continuously beginning very early after hatching. Proceedings of the National Academy of Sciences 119(18):e2122052119. 10.1073/pnas.2122052119

Finenko, G.A., Kideys, A.E., Anninsky, B.E., Shiganova, T.A., Roohi, A., Tabari, M.R., Rostami, H., Bagheri, S. (2006) Invasive ctenophore *Mnemiopsis leidyi* in the Caspian Sea: feeding, respiration, reproduction and predatory impact on the zooplankton community. Marine Ecology Progress Series 314:171–185. 10.3354/meps314171

Finenko, G.A., Pavlovskaya, T.V., Romanova, Z.A., Abolmasova, G I., Datsyk, N.A. (2008) Digestion time and feeding intensity of the larval ctenophore *Mnemiopsis leidyi* A. Agassiz (Ctenophora, Lobata). Marine Ecological Journal 7(3):61– 74.

Firth, D. (1993) Bias reduction of maximum likelihood estimates. Biometrika 80:27–38. 10.2307/2336755

Garbe, A. (1901) Untersuchungen über die Entstehung der Geschlechtsorgane bei den Ctenophoren. Zeitschrift für wissenschaftliche Zoologie 69:472–491

Granhag, L., Møller, L.F., Hansson, L.J. (2011) Size-specific clearance rates of the ctenophore *Mnemiopsis leidyi* based on in situ gut content analyses. Journal of Plankton Research 33(7):1043–1052. 10.1093/plankt/fbr010

Granhag, L., Hosia, A. (2015) Feeding and starvation in the native ctenophore *Bolinopsis infundibulum* and the introduced *Mnemiopsis leidyi* in the North Sea: implications for ctenophore transport in ships’ ballast water. Journal of Plankton Research 37(5):1006–1010. 10.1093/plankt/fbv058

Harris, R.N. (1987) Density-dependent paedomorphosis in the salamander *Notophthalmus viridescens dorsalis*. Ecology 68(3):705–712. 10.2307/1938476

Hirota, J. (1972) Laboratory culture and metabolism of the planktonic ctenophore, Pleurobrachia bachei A. Agassiz. In: Motoda, S., Takenouti, Y.A. (eds) Biological Oceanography of the Northern North Pacific Ocean. Idemitsu Shoten, Tokyo, pp 465–484

Hosia, A., Falkenhaug, T. (2015) Invasive ctenophore *Mnemiopsis leidyi* in Norway. Marine Biodiversity Records 8:e31. 10.1017/S1755267215000044

Jaspers, C., Møller, L.F., Kiørboe, T. (2011) Salinity gradient of the Baltic Sea limits the reproduction and population expansion of the newly invaded comb jelly *Mnemiopsis leidyi*. PLoS ONE 6(8):e24065. 10.1371/journal.pone.0024065

Jaspers, C., Haraldsson, M., Bolte, S., Reusch, T.B.H., Thygesen, U.H., Kiørboe, T. (2012) Ctenophore population recruits entirely through larval reproduction in the central Baltic Sea. Biology Letters 8(5):809–812. 10.1098/rsbl.2012.0163

Jaspers, C., Costello, J.H., Colin, S.P. (2014) Carbon content of *Mnemiopsis leidyi* eggs and specific egg production rates in northern Europe. Journal of Plankton Research 37(1):11–15. 10.1093/plankt/fbu102

Jaspers, C., Møller, L.F., Kiørboe, T. (2015) Reproduction rates under variable food conditions and starvation in *Mnemiopsis leidyi*: significance for the invasion success of a ctenophore. Journal of Plankton Research 37(5):1011–1018. 10.1093/plankt/fbv017

Jaspers, C., Marty, L., Kiørboe, T. (2017) Selection for life-history traits to maximize population growth in an invasive marine species. Global Change Biology 24(3):1164–1174. 10.1111/gcb.13955

Javidpour, J., Molinero, J.C., Ramírez-Romero, E., Roberts, P., Larsen, T. (2020) Cannibalism makes invasive comb jelly, Mnemiopsis leidyi, resilient to unfavourable conditions. Communications Biology 3:212. 10.1038/s42003-020-0940-2

Kideys, A.E. (2002) Fall and rise of the Black Sea ecosystem. Science 297(5586):1482–1484. 10.1126/science.1073002

Kim, I.V., Navarrete, C., Grau-Bové, X., Iglesias, M., Elek, A., Zolotarov, G., Bykov, N.S., Montgomery, S.A., Ksiezopolska, E., Cañas-Armenteros, D., Soto-Angel, J.J., Leys, S.P., Burkhardt, P., Suga, H., de Mendoza, A., Marti-Renom, M.A., Sebé-Pedrós, A. (2025) Chromatin loops are an ancestral hallmark of the animal regulatory genome. Nature 642(8069):1097– 1105. 10.1038/s41586-025-08960-w

Koutsouveli, V., Torres-Oliva, M., Bayer, T., Fuß, J., Grossschmidt, N., Marulanda-Gomez, A.M., Jensen, N., Gill, D., Schmitz, R.A., Pita, L., Reusch, T.B.H. (2025) The chromosome-level genome of the ctenophore *Mnemiopsis leidyi* A. Agassiz, 1865 reveals a unique immune gene repertoire. Genome Biology and Evolution 17(2):evaf006. 10.1093/gbe/evaf006

Kremer, P.M., 1976. The ecology of the ctenophore Mnemiopsis leidyi in Narragansett Bay. Ph.D. Thesis, University of Rhode Island: 325 pp. 10.23860/diss-2825

Lenth, R.V., Piaskowski, J. (2026) emmeans: Estimated Marginal Means, aka Least-Squares Means. R package

Malej, A., Lučić, D., Tirelli, V., Vodopivec, M., Bettoso, N., Ferrari, C.R., Camatti, E., Paliaga, P., Shiganova, T. (2017) Invasive *Mnemiopsis leidyi* (Ctenophora) in the Northern Adriatic: here to stay? ASLO 2017 Conference – Mountains to the Seas. Honolulu, Hawaii.

Marshall, D.J., Keough, M.J. (2008) The evolutionary ecology of offspring size in marine invertebrates. Advances in Marine Biology 53:1–60. 10.1016/S0065-2881(07)53001-4

Martindale, M.Q. (1987) Larval reproduction in the ctenophore *Mnemiopsis mccradyi* (order Lobata). Marine Biology 94(3):409–414. 10.1007/BF00428247

Mugabo, M., Perret, S., Legendre, S., Le Galliard, J.-F. (2013) Density-dependent life history and the dynamics of small populations. Journal of Animal Ecology 82(6):1227–1239. 10.1111/1365-2656.12109

Müller, T., Küll, C.L. Müller, C. (2016) Effects of larval versus adult density conditions on reproduction and behavior of a leaf beetle. Behavioral Ecology and Sociobiology 70:2081–2091. 10.1007/s00265-016-2212-1

Oguz, T., Fach, B., Salihoglu, B. (2008) Invasion dynamics of the alien ctenophore *Mnemiopsis leidyi* and its impact on anchovy collapse in the Black Sea. Journal of Plankton Research 30(12):1385–1397. 10.1093/plankt/fbn094

Patry, W.L., Bubel, M., Hansen, C., Knowles, T. (2020) Diffusion tubes: a method for the mass culture of ctenophores and other pelagic marine invertebrates. PeerJ 8:e8938. 10.7717/peerj.8938

Pianka, E.R. (1970) On r- and K-selection. The American Naturalist 104(940):592–597. 10.1086/282697

Piccardi, F., Poli, F., Sguotti, C., Tirelli, V., Borme, D., Mazzoldi, C., Barausse, A. (2025) Assessing the impact of the invasive ctenophore *Mnemiopsis leidyi* on artisanal fisheries in the Venice Lagoon: an interdisciplinary approach. Hydrobiologia 852(8–9):2387–2405. 10.1007/s10750-024-05505-6

Pinheiro, J., Bates, D., R Core Team (2024) nlme: Linear and Nonlinear Mixed Effects Models. R package

Presnell, J.S., Bubel, M., Knowles, T., Patry, W., Browne, W.E. (2022) Multigenerational laboratory culture of pelagic ctenophores and CRISPR–Cas9 genome editing in the lobate *Mnemiopsis leidyi*. Nature Protocols 17(8):1868–1900. 10.1038/s41596-022-00702-w

Purcell, J.E., Shiganova, T.A., Decker, M.B., Houde, E.D. (2001) The ctenophore *Mnemiopsis* in native and exotic habitats: U.S. estuaries versus the Black Sea basin. Hydrobiologia 451:145–176. 10.1023/A:1011826618539

Ramon-Mateu, J., Ellison, S.T., Angelini, T.E., Martindale, M.Q. (2019) Regeneration in the ctenophore *Mnemiopsis leidyi* occurs in the absence of a blastema, requires cell division, and is temporally separable from wound healing. BMC Biology 17(1):80. 10.1186/s12915-019-0695-8

R Core Team (2024) R: A Language and Environment for Statistical Computing. R Foundation for Statistical Computing, Vienna, Austria

Rathjen, K.A., Breitburg, D.L., Neale, P.J. (2012) Effects of ultraviolet radiation on the growth, reproduction and survival of the lobate ctenophore *Mnemiopsis leidyi* in Chesapeake Bay. Journal of Experimental Marine Biology and Ecology 432–433:121–130. 10.1016/j.jembe.2012.06.029

Rečnik, K., Klun, K., Lipej, L., Malej, A., Tinta, T. (2024) Chemical composition and egg production capacity throughout bloom development of ctenophore *Mnemiopsis leidyi* in the northern Adriatic Sea. PeerJ 12:e17844. 10.7717/peerj.17844

Reeve, M.R., Walter, M.A. (1978) Nutritional ecology of ctenophores — a review of recent research. In: Russell, F.S. et al (eds) Advances in Marine Biology, vol 15. Academic Press, London, pp 249–287

Reichert, S., Berger, V., Jackson, J., Chapman, S.N., Htut, W., Mar, K.U., Lummaa, V. (2019) Maternal age at birth shapes offspring life-history trajectory across generations in long-lived Asian elephants. Journal of animal ecology 89(4):996–1007. 10.1111/1365-2656.13049

Reznick, D.A., Bryga, H., Endler, J.A. (1990) Experimentally induced life-history evolution in a natural population. Nature 346:357–359. 10.1038/346357a0

Richter, J., Martin, L., Beachy, C.K. (2009) Increased larval density induces accelerated metamorphosis independently of growth rate in the frog *Rana sphenocephala*. Journal of Herpetology 43(3):551–554. 10.1670/08-060R1.1

Riisgård, H.U., Bøttiger, L., Madsen, C.V., Purcell, J.E. (2007) Invasive ctenophore *Mnemiopsis leidyi* in Limfjorden (Denmark) in late summer 2007-assessment of abundance and predation effects. Aquatic Invasions 2(4):395–401. 10.3391/ai.2007.2.4.8

Ryan, J.F., Pang, K., Schnitzler, C.E., Nguyen, A.D., Moreland, R.T., Simmons, D.K., Koch, B.J., Francis, W.R., Havlak, P., NISC Comparative Sequencing Program, Smith, S.A., Putnam, N.H., Haddock, S.H.D., Dunn, C.W., Wolfsberg, T.G., Mullikin, J.C., Martindale, M.Q., Baxevanis, A.D. (2013) The genome of the ctenophore Mnemiopsis leidyi and its implications for cell type evolution. Science 342(6164):1242592. 10.1126/science.1242592

Sasson, D.A., Ryan, J.F. (2016) The sex lives of ctenophores: the influence of light, body size, and self-fertilization on the reproductive output of the sea walnut, *Mnemiopsis leidyi*. PeerJ 4:e1846. 10.7717/peerj.1846

Sasson, D.A., Jacquez, A.A., Ryan, J.F. (2018) The ctenophore *Mnemiopsis leidyi* regulates egg production via conspecific communication. BMC Ecology 18:12. 10.1186/s12898-018-0169-9

Schnitzler, C.E., Pang, K., Powers, M.L., Reitzel, A.M., Ryan, J.F., Simmons, D., Tada, T., Park, M., Gupta, J., Brooks, S.Y., Blakesley, R.W., Yokoyama, S., Haddock, S.H.D., Martindale, M.Q., Baxevanis, A.D. (2012) Genomic organization, evolution, and expression of photoprotein and opsin genes in *Mnemiopsis leidyi*: a new view of ctenophore photocytes. BMC Biology 10:107. 10.1186/1741-7007-10-107

Sebé-Pedrós, A., Chomsky, E., Pang, K., Lara-Astiaso, D., Gaiti, F., Mukamel, Z., Amit, I., Hejnol, A., Degnan, B.M., Tanay, A. (2018) Early metazoan cell type diversity and the evolution of multicellular gene regulation. Nature Ecology & Evolution 2(7):1176–1188. 10.1038/s41559-018-0575-6

Shiganova, T.A., Bulgakova, Y.V. (2000) Effects of gelatinous plankton on Black Sea and Sea of Azov fish and their food resources. ICES Journal of Marine Science 57(3):641–648. 10.1006/jmsc.2000.0736

Shiganova, T.A. (2020) Adaptive strategies of *Mnemiopsis leidyi* A. Agassiz 1865 in different environments of the Eurasian seas. Marine Pollution Bulletin 161:111737. 10.1016/j.marpolbul.2020.111737

Siapatis, A., Giannoulaki, M., Valavanis, V.D., Palialexis, A., Schismenou, E., Machias, A., Somarakis, S. (2008) Modelling potential habitat of the invasive ctenophore *Mnemiopsis leidyi* in Aegean Sea. Hydrobiologia 612(1):281–295. 10.1007/s10750-008-9497-7

Silakov, M.I., Kuznetsov, A.V., Temnykh, A.V., Anninsky, B.E. (2023) Effect of monochromatic light on the behavior of the ctenophore *Mnemiopsis leidyi*. Biosystems 231:104987. 10.1016/j.biosystems.2023.104987

Sobrero, L., Chiaverano, L.M., Macchi, G.J., Schiariti, A. (2026) Temperature sets a lower threshold for reproduction but has no detectable effect on reproductive output in *Mnemiopsis leidyi* (Ctenophora). Marine Biology 173(10):169. 10.1007/s00227-026-04928-0

Soto-Angel, J.J., Jaspers, C., Hosia, A., Majaneva, S., Martell, L., Burkhardt, P. (2023) Are we there yet to eliminate the terms larva, metamorphosis, and dissogeny from the ctenophore literature? Proceedings of the National Academy of Sciences 120(4):e2218317120. 10.1073/pnas.2218317120

Soto-Angel, J.J., Nordmann, E.L., Sturm, D., Sachkova, M., Pang, K., Burkhardt, P. (2024) Stable laboratory culture system for the ctenophore *Mnemiopsis leidyi*. In: Moroz, L.L. (ed) Ctenophores, Methods in Molecular Biology, vol 2757. Humana, New York, pp 123–145. 10.1007/978-1-0716-3642-8_4

Soto-Angel, J.J., Burkhardt, P. (2024) Reverse development in the ctenophore *Mnemiopsis leidyi*. Proceedings of the National Academy of Sciences 121(45):e2411499121. 10.1073/pnas.2411499121

Stelzer, C.P., Snell, T.W. (2003) Induction of sexual reproduction in *Brachionus plicatilis* (Monogononta, Rotifera) by a density-dependent chemical cue. Limnology and Oceanography 48(2):939–943. 10.4319/lo.2003.48.2.0939

Sullivan, L.J. (2010). Gut evacuation of larval *Mnemiopsis leidyi* A. Agassiz (Ctenophora, Lobata). Journal of Plankton Research 32(1):69–74. 10.1093/plankt/fbp100

Therneau, T.M. (2024) A Package for Survival Analysis in R. R package

Therneau, T.M., Grambsch, P.M. (2000) Modeling Survival Data: Extending the Cox Model. Springer, New York

Totton, A.K. (1954) Egg-laying in Ctenophora. Nature 174(4425):360. 10.1038/174360a0

Venables, W.N., Ripley, B.D. (2002) Modern Applied Statistics with S, 4th ed. Springer, New York

Wang, S., Zhang, G., Zhou, K., Sun, S. (2020) Long-term population variability and reproductive strategy of a northward expanded ctenophore *Pleurobrachia globosa* Moser, 1903 in a temperate bay, China. Journal of Experimental Marine Biology and Ecology 533:151457. 10.1016/j.jembe.2020.151457

Wilbur, H.M., Collins, J.P. (1973) Ecological aspects of amphibian metamorphosis: nonnormal distributions of competitive ability reflect selection for facultative metamorphosis. Science 182(4119):1305–1314. 10.1126/science.182.4119.1305

Wong, S.S., Yu, J., Schroeder, F.C., Kim, D.H. (2020) Population density modulates the duration of reproduction of *C. elegans*. Current Biology 30(13):2602–2607.e2. 10.1016/j.cub.2020.04.056

Wood, S.N. (2011) Fast stable restricted maximum likelihood and marginal likelihood estimation of semiparametric generalized linear models. Journal of the Royal Statistical Society (B) 73:3–36. 10.1111/j.1467-9868.2010.00749.x

Wood, S.N. (2017) Generalized Additive Models: An Introduction with R, 2nd ed. Chapman and Hall/CRC

Yazdani Foshtomi, M., Abtahi, B., Esmaili Sari, A., Taheri, M. (2007) Ion composition and osmolarity of Caspian Sea ctenophore, *Mnemiopsis leidyi*, in different salinities. Journal of Experimental Marine Biology and Ecology 352(1):28–34. 10.1016/j.jembe.2007.06.029

